# Complex spikes perturb movements, revealing the sensorimotor map of Purkinje cells

**DOI:** 10.1101/2023.04.16.537034

**Authors:** Salomon Z. Muller, Jay S. Pi, Paul Hage, Mohammad Amin Fakharian, Ehsan Sedaghat-Nejad, Reza Shadmehr

**Affiliations:** Laboratory for Computational Motor Control, Dept. of Biomedical Engineering, Johns Hopkins School of Medicine, Baltimore, Maryland USA; Zuckerman Mind Brain Behavior Institute, Department of Neuroscience, Columbia University, New York, NY USA

## Abstract

The cerebellar cortex performs computations that are critical for control of our actions, and then transmits that information via simple spikes of Purkinje cells (P-cells) to downstream structures. However, because P-cells are many synapses away from muscles, we do not know how their output affects behavior. Furthermore, we do not know the level of abstraction, i.e., the coordinate system of the P-cell’s output. Here, we recorded spiking activities of hundreds of P-cells in the oculomotor vermis of marmosets during saccadic eye movements and found that following the presentation of a visual stimulus, the olivary input to a P-cell encoded a probabilistic signal that coarsely described both the direction and the amplitude of that stimulus. When this input was present, the resulting complex spike briefly suppressed the P-cell’s simple spikes, disrupting the P-cell’s output during that saccade. Remarkably, this brief suppression altered the saccade’s trajectory by pulling the eyes toward the part of the visual space that was preferentially encoded by the olivary input to that P-cell. Thus, analysis of behavior in the milliseconds following a complex spike unmasked how the P-cell’s output influenced behavior: the preferred location in the coordinates of the visual system as conveyed probabilistically from the inferior olive to a P-cell defined the action in the coordinates of the motor system for which that P-cell’s simple spikes directed behavior.

**Significance:** We are lacking general principles that can describe how changes in a P-cell’s simple spikes might alter behavior. Here, we show that a brief suppression of a P-cell’s simple spikes in the oculomotor vermis consistently pulls the eyes in a direction that corresponds to the preferred location of the sensory space as conveyed probabilistically to that P-cell from the inferior olive. Thus, the inferior olive defines the coordinate system regarding the information that a P-cell is providing to the rest of the brain.

## Introduction

To understand the computations that are performed by a neuron, it is useful to quantify how the change in its output is translated into changes in behavior. In practice this is challenging because in the brain, neurons are often many synapses away from muscles. This distance not only presents a connectome challenge, namely, the difficulty to trace from a muscle back to a neuron, but more importantly a challenge of representation: as signals pass through the many layers of neurons, information converges from various areas of the brain and undergoes transformations before a motor command to a muscle is generated. For example, in the cerebellum, a network of neurons in the cerebellar cortex may make predictions that are critical for control of a behavior, which are then transmitted to the cerebellar nucleus via the simple spikes of Purkinje cells (P-cells). But what is the contribution of a P-cell’s simple spikes to the control of that behavior?

We may probe the behavioral effects of a group of neurons through lesion experiments, or via optogenetics, silencing or activating cells to study their downstream behavioral effects. In cerebellum, however, we have a unique opportunity: a P-cell conveys its output via simple spikes (SSs) at a rate of 50-100 Hz, and receives a single climbing fiber input that produces a complex spike (CS) at around 1 Hz. Critically, the CS robustly but briefly suppresses the SS (1, 2). Does this suppression affect behavior? If so, what does the behavioral change imply regarding the information that is normally transmitted by that P-cell?

Anatomical and stimulation studies have hinted that the climbing fiber input of a P-cell likely plays a central role in defining the downstream projections of that P-cell (3, 4). This input may encode sensory events (5) and transmit it to a handful of P-cells, which anatomically converge onto the same or nearby nucleus neurons (6–8). Stimulation of that region of the cerebellar nucleus produces a movement that appears related to the climbing fiber input to the P-cells (9). For example, touching the forelimb results in complex spikes (CSs) in a group of P-cells that converge onto a region in the interposed nucleus, and stimulation of that part of the interposed nucleus results in the activation of muscles that withdraw the forelimb (9). Optogenetic activation of P-cells in a region of the oculomotor vermis that receives climbing fiber inputs that mostly encode for visual events to the right of the fovea produce movements of the eyes toward the left (10). Thus, the encoding of the sensory space by the climbing fiber input to a P-cell may play a critical role in defining the downstream effects that the P-cell’s simple spikes (SSs) have on behavior.

To examine this idea, here we relied on the fact that a P-cell receives only a single climbing fiber, producing a CS that not only provides information about the sensory event (5, 11), but also briefly suppresses the SSs. In the oculomotor region of the vermis, each climbing fiber preferentially encodes a particular part of the visual space, but the SSs are modulated for all saccades (12, 13). Occasionally, a CS occurs before a saccade, suppressing the SSs that the P-cell would normally produce during that movement (14, 15). This allowed us to quantify the relationship between the sensory encoding of the visual space in the climbing fiber input to a P-cell, and the motor consequences of that P-cell’s SS suppression. The resulting sensorimotor map revealed an alignment: the preferred region of the sensory space, as encoded by the climbing fiber input to a P-cell, was aligned with the direction of force production that followed the suppression of the SSs of that P-cell.

## Results

To quantify the encoding of the visual space by the climbing fiber input to a P-cell, we trained marmosets to fixate a central location and presented a primary target at one of eight random directions (Fig. 1A-B). As the primary saccade commenced, we moved the target to another random location, instructing the subject to make a secondary saccade (Fig. 1A-B). Following 200 ms fixation, the center target was re-displayed, and the subject made a center saccade to begin another trial.

**Figure 1.**
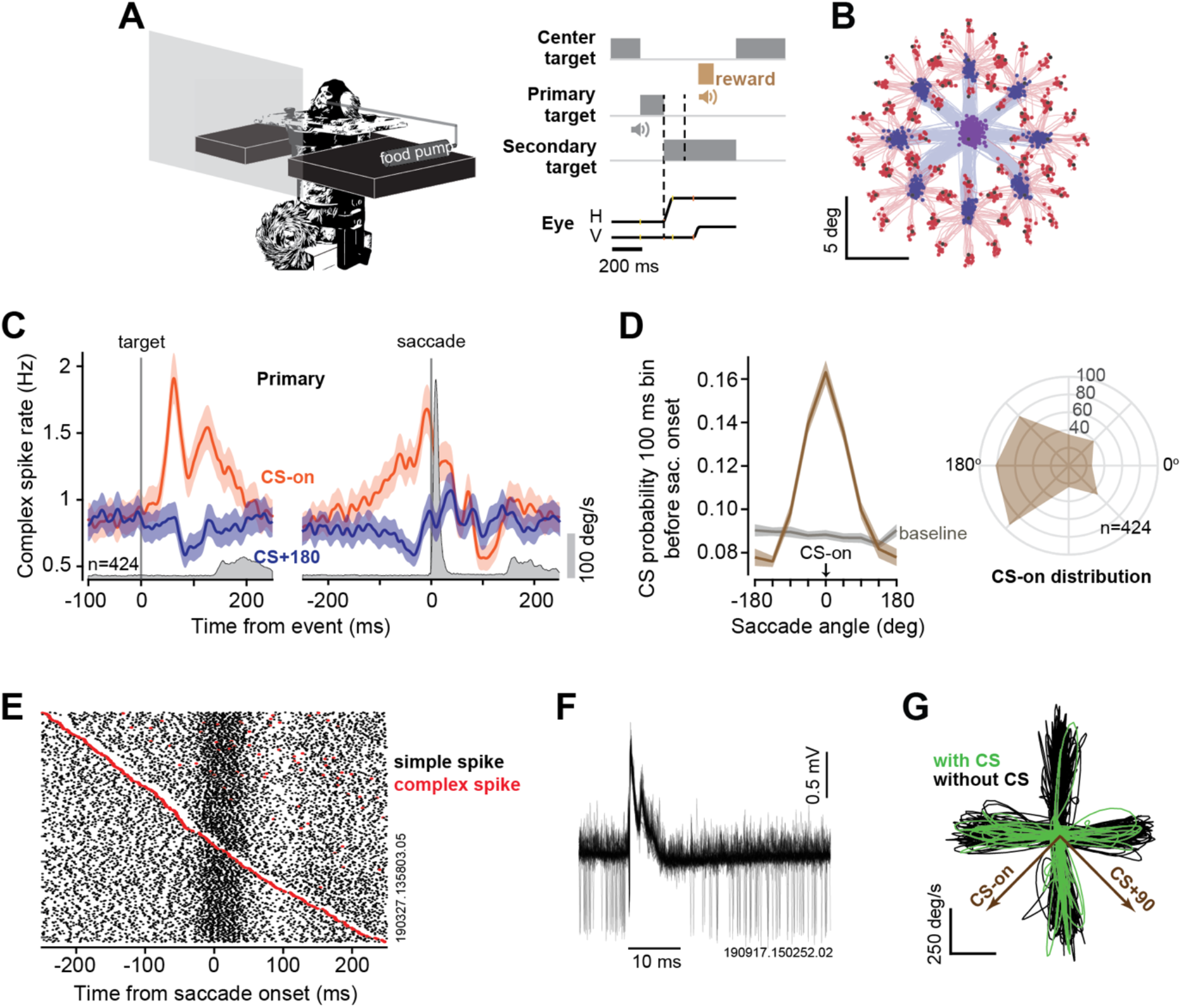
Complex spikes of P-cells in the oculomotor region of the cerebellar vermis responded to target presentation as well as saccades. **A**. Experimental paradigm. Marmosets fixated a center target and then made a saccade to a primary target placed randomly at one of 8 directions. During the primary saccade the target was moved to another random location, resulting in a secondary saccade. **B**. Eye position traces for the primary (blue) and secondary (red) saccades. **C**. CS response aligned to the onset of the primary target, and the onset of the saccade. Gray trace is eye velocity. CS-on was defined as the target direction that during the 200 ms after target onset produced the largest increase in CS rates (see Methods). **D**. Left plot: CS probability in the 100 ms bin before saccade onset with respect to CS-on, demonstrating that many of the CSs occurred just before saccade onset. Right: the distribution of CS-on directions across the P-cells. **E**. Simple spike rasters were aligned on saccade onset and ordered by timing of the CS. This P-cell exhibited a burst of SS activity near saccade onset, but occurrence of a CS briefly suppressed that modulation. **F**. Extracellular recording from an example P-cell: the CS suppresses the SS for about 10 ms. **G**. Trajectories in velocity space for a subset of the primary saccades (4 directions are shown). Some saccades were preceded by a complex spike (green). The CS-on vector indicates the target direction that produced the largest increase in CS rates.

We recorded activities of n=424 P-cells in lobules VI and VII of the cerebellar vermis in 162 recording sessions over the course of 2.5 years in two marmosets (291 P-cells in subject M, 133 P-cells in subject R). In every case, the neuron was identified as a P-cell because of the presence of CSs. In n=281 P-cells we were able to isolate both the CSs and the SSs and verified that the CS suppressed the SS. In 59 recording sessions we were able to simultaneously isolate two or more P-cells, allowing us to examine the effects of simultaneous CSs on behavior.

Presentation of the target modulated the CS firing rates (Fig. 1C). For each P-cell we estimated the target direction that produced the largest increase in CS rates (Methods) and labeled it as the CS-on direction (Fig. 1C-D). As a result, the largest CS rates were in response to a target in direction CS-on, and the lowest response was, on average, in direction CS+180 (baseline rates were defined during the fixation period) (Fig. 1D, left subplot). As many of our recordings were from the right side of the vermis (152 P-cells recorded from the right vermis, 217 from the medial vermis, 55 from the left vermis), the distribution of CS-on directions across the P-cells was biased toward the left side (12, 14) (Fig. 1D, right subplot).

Sometimes the CS occurred just before the onset of a saccade (Fig. 1E). This briefly suppressed the simple spikes that the P-cell would normally produce during that movement (Fig. 1E-F). To quantify the effects of SS suppression on behavior, we compared saccade trajectories that were made toward a given target with and without a CS. That is, we compared saccades for which at least one CS occurred during the −90 to +10 ms period with respect to movement onset (Fig. 1G, with CS), with saccades that were free of a CS during the same period (Fig. 1G, without CS). We focused on the saccades that were made from center to the primary target because these saccades started from the same location and were in response to the same target.

As expected, if a CS occurred in the −90 to +10 ms period with respect to saccade onset, then the SS production of the P-cell was suppressed (Fig. 2A, left subplot). In this figure, the SS suppression is not to zero because the CS could occur at any time during the 100 ms period. As a result, pooling across all CS events muted the resulting average SS suppression, but as confirmation we aligned the data to the onset of the CS, and indeed found that the SS production was suppressed (Fig. 2A, right subplot).

**Figure 2.**
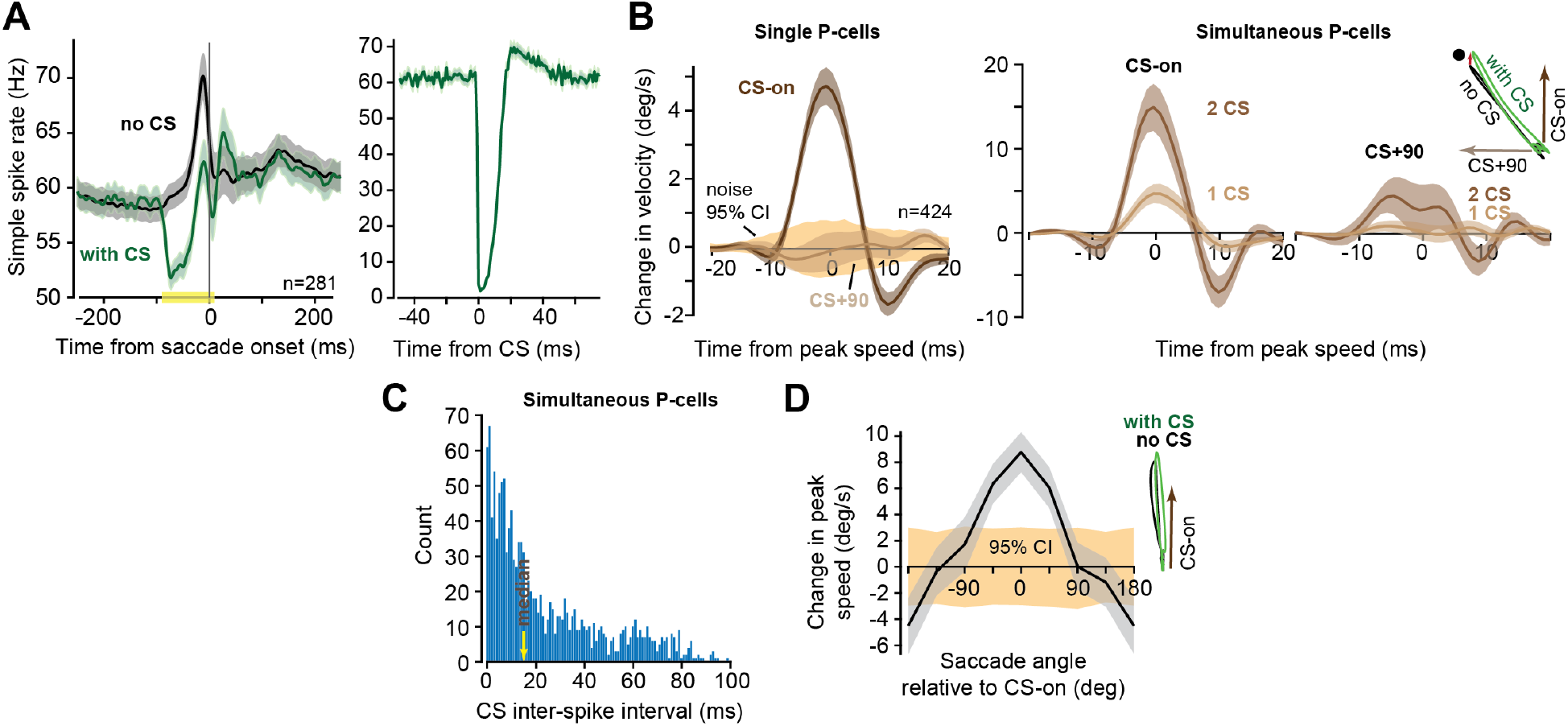
The CS suppressed SS production of the P-cell, pulling the eyes in direction CS-on. **A**. Left: for each P-cell, the SS activity was aligned to saccade onset and grouped based on whether the P-cell had a CS in the 100 ms interval around saccade onset (yellow marker) or not. Right plot: same data aligned to CS onset. **B**. Left: effect of CS on saccade trajectory (sequence of velocity vectors), as projected onto the CS-on and CS+90 vectors of each P-cell. 95% CI refers to the bootstrap estimate of the confidence interval. The effect of a CS was to pull the eyes in direction CS-on. Right: data from the subset of recordings in which two or more P-cells were isolated and either one or two CSs were present in the −90 to +10 ms period around saccade onset. **C**. Distribution of inter-spike intervals between the two CSs that were recorded in the simultaneous P-cells. **D**. Effects of CS on saccade peak velocity. When the target was in direction CS-on of the P-cell, a CS resulted in a slightly faster movement. When the target was in direction CS+180, a CS resulted in a slightly slower movement. Thus, regardless of saccade direction, a CS pulled the eyes in direction CS-on. Error bars are SEM. 95% CI refers to the bootstrap estimate of the confidence interval for the effects of random splitting of the two groups on changes in saccade peak speed. Note that for this and similar plots in this paper, the value for −180 angle is repeated for the 180 angle for symmetrical viewing. Error bars are SEM or 95% CI.

### Suppression of the simple spikes pulled the eyes in direction CS-on

We began by quantifying the effects that the CS of a single P-cell had on behavior. We separated saccades into those that were preceded by a CS, and those that were not. We represented a saccade’s trajectory as a time sequence of velocity vectors and then computed average of this time sequence across all with-CS saccades, minus the time sequence averaged across all without-CS saccades, for each target. This produced a sequence of vectors that described the CS induced change in velocity at each instant of time during the movement to a target. To test the hypothesis that the sensory CS-on space defines the motor output, we then projected this sequence of vectors onto an orthonormal coordinate system: the CS-on and the CS+90 vectors of that P-cell. The resulting projection produced two scalar quantities as a function of time: one quantity measured the SS suppression induced deviation of the saccade in direction CS-on, and then other quantity measured the deviation in direction CS+90. Finally, we averaged each of these two quantities across target directions for that P-cell, and then across all P-cells. We found that the SS suppression had a behavioral consequence, producing a deviation that was, on average, entirely in direction CS-on, as there were no changes in direction CS+90 (Fig. 2B, left subplot).

The peak of the trajectory perturbation was only about 5 deg/s, which is two orders of magnitude smaller than the peak speed of the saccade (Fig. 1G), around 1% of the movement itself. This appears reasonable because the SS suppression that a CS causes is so short-lived (around 10 ms) that typically, it would remove only one or two SS spikes that would have been produced by the P-cell during the movement. To test for statistical reliability, we performed a bootstrapping procedure in which for each P-cell, and for each direction, we randomly split the saccades into two groups (rather than based on the occurrence of a CS) while keeping the ratio of saccade numbers as in the non-random split. We then projected the difference between the two groups of saccades onto the CS-on of that P-cell and from the resulting distribution we computed the 95% confidence interval (CI). This revealed that if a CS was present around saccade onset, then the saccade’s trajectory was deviated in direction CS-on by an amount that was 4.5 times larger than the 95% CI (Fig. 2B, left subplot).

### Multiple complex spikes in nearby P-cells magnified the perturbation

We next focused on the 59 sessions in which we were able to simultaneously isolate two or more P-cells. In each dataset we selected only the P-cells that had a similar CS-on, i.e., the P-cells in which the CS-on directions fell within the same 45 deg bin (see Methods for a breakdown of the number of P-cells in the various sessions). We then considered saccades in which none of the simultaneously recorded P-cells had a CS, saccades in which only one P-cell had a CS, and saccades in which two or more P-cells had a CS in the −90 to +10 ms period before saccade onset. This allowed us to quantify the behavioral effects of multiple CSs.

We found that the presence of two CSs dramatically increased the size of the trajectory deviation, more than tripling the pull in direction CS-on, as compared to a single CS (Fig. 2B, right subplot). Notably, the effect of multiple CSs on the saccade’s trajectory in direction CS+90 remained smaller than in direction CS-on (whereby projection in direction CS+90 was within the 95% CI, Fig. S1A, left subplot). The CS inter-spike interval (ISI) in the simultaneously recorded P-cells had an exponential distribution with a median of 15 ms (Fig. 2C), implying that most of the CS events were synchronous. However, splitting the data based on CS ISI did not significantly affect the results (Fig. S1A, right subplot).

In summary, the presence of a CS near saccade onset altered the saccade’s trajectory, pulling the eyes in direction CS-on of the P-cell. This effect more than tripled if the CSs were present in two or more simultaneously recorded P-cells that had a similar CS-on.

### The CS-induced effect depended on the direction of the movement

According to our framework, the effect of a CS on the saccade should depend on the direction of the movement. For example, if the target is in direction CS-on of the P-cell, then the CS induced SS suppression in that P-cell should pull the eyes toward that target. This should produce a slightly greater peak speed than saccades that are made toward the same target but without a preceding CS. In contrast, if the target happens to be in direction CS+180 of the P-cell, then the presence of a CS in that P-cell should pull the eyes in the opposite direction, thus slowing the saccade.

To test these predictions, we separated the saccades based on the direction of the target with respect to CS-on. We observed that when a saccade was toward a target in direction CS-on of the P-cell, then that saccade tended to have a greater peak speed if the CS was present before saccade onset as compared to when it was absent. In contrast, if the saccade was toward a target in direction CS+180, presence of a CS produced a saccade that tended to have a lower peak speed (peak-speed difference between CS-on and CS+180 is ∼13 deg/s, both values are with p<0.01, Fig. 2D). Thus, regardless of movement direction, the CS and the accompanying SS suppression pulled the eyes in direction CS-on of the P-cell.

### Control studies

If the SS suppression is necessary for pulling the eyes in direction CS-on, then the CS that occurs before a saccade but does not disrupt the saccade-related SS modulation should not affect the saccade’s trajectory. To check for this, we considered CSs that occurred in the period just before the onset of SS modulation, i.e., the period −190 to −90 ms before saccade onset. These early CSs suppressed the SSs that were present during fixation, but not the SSs that were present before saccade onset (Fig. 3A, left subplot). The saccades that followed these early CSs did not show a robust change in any direction (Fig. 3B, values are within 95%CI of bootstrap analysis). Similarly, CSs that occurred late in the saccade’s trajectory (Fig. 3A right subplot, +10 to +40 ms after saccade onset) had no effect on the saccade (Fig. 3B).

**Figure 3.**
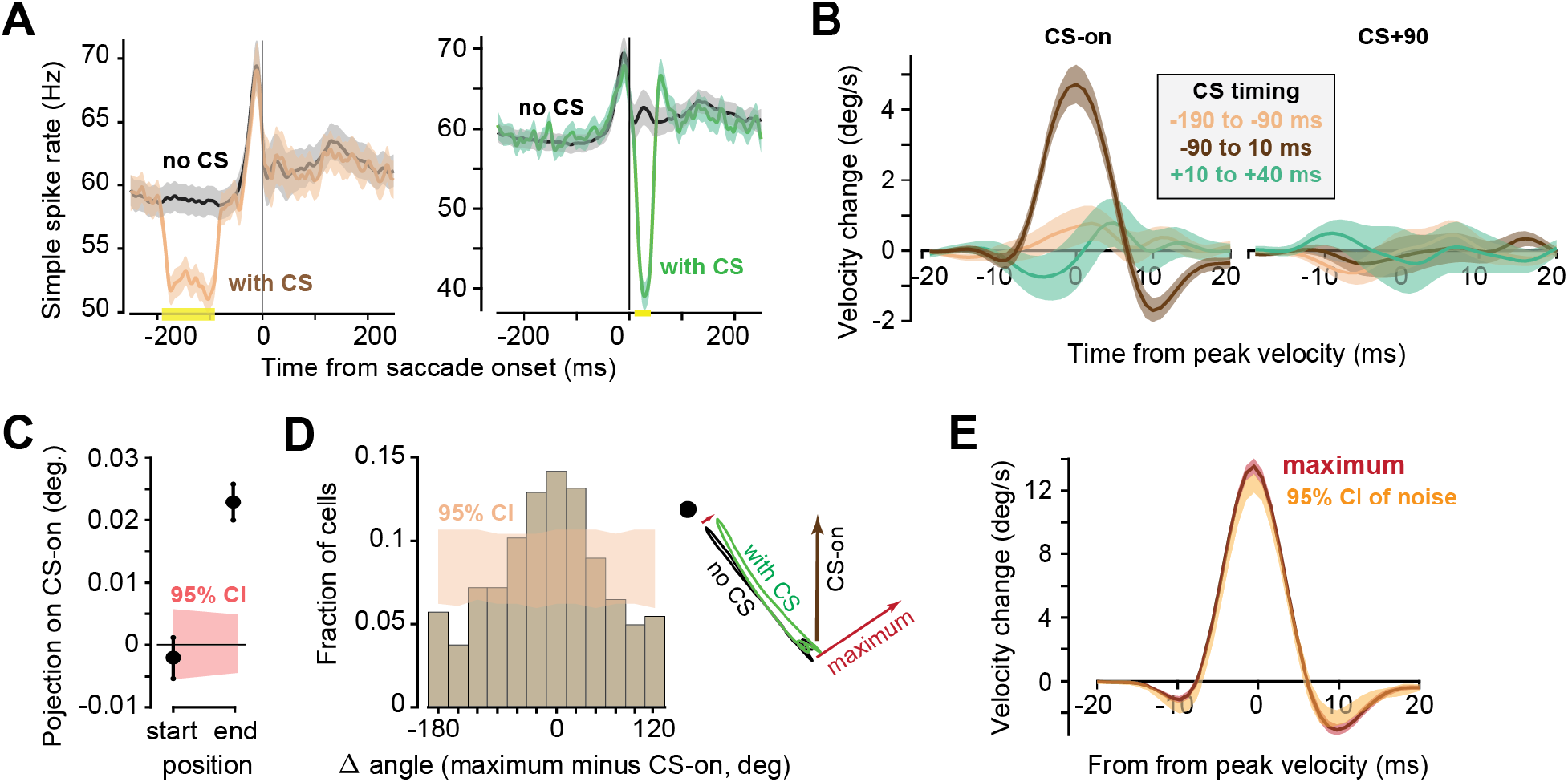
CS-on vector is an unbiased estimator of the direction of perturbation. **A**. Left: the effect of a CS that occurred −90 to −190 ms before saccade onset on simple spikes. Right: the effect a CS that occurred +10 to +40 ms after saccade onset on simple spikes. **B**. The effect of the timing of the CS on saccade trajectory. Neither the early nor the late CSs had a significant effect on the saccade. **C.** Primary saccade starting points do not significantly differ between with- and without-CS saccades when projected on the CS-on vector. Rather, the final points significantly differ suggesting that the CS perturbs the movement in direction CS-on. **D**. Angular distance between the CS-on direction of a P-cell, and the unit vector that maximized the projection of the CS-induced change in saccade trajectory (that unit vector was labeled as the “maximum” vector). As an example, the right plot shows trajectories of two saccades in velocity space, and the maximum vector. **E**. Projection of the CS-induced changed in saccade trajectory onto the maximum vector. The 95% CI indicates the effect size due to random noise. Compare the 95% CI here with Fig. 2B. Thus, CS-on is both an unbiased and a low-noise estimate of the direction of CS-induced perturbation. Error bars are SEM or 95% CI.

Our comparison of saccades with and without a CS required that the starting point of the two saccades be the same. For example, if the with-CS saccades were, for some reason, consistently biased so that the start point was toward direction CS+180, which may slightly increase the probability of firing a CS, then the effects that we have documented could not be attributed to a movement disruption. To check for this, we measured the difference in the starting points of the two groups of saccades and projected that difference on the CS-on vector. The difference in the starting points of the with-CS and without-CS saccades was near zero and well within 95% CI (Figure 3C). Difference in the final points, on the other hand, were significantly in direction CS-on (Figure 3C, p<0.001 as measure by a bootstrapping procedure). Furthermore, analyzing differences in the saccade offset and saccade amplitude of with- versus without-CS-saccades while normalizing to the different starting points (see Methods for details of the analysis) showed that the effects on amplitude and saccade endpoint remained robust to the small changes in saccade starting position (Fig. S1B).

### Is the CS-on vector a good estimate of the direction of perturbation?

We quantified the behavioral effects by projecting the saccade’s trajectory onto a specific coordinate system: the CS-on of that P-cell. However, we do not know if this is the best coordinate system that we could have chosen. To answer this question, for each P-cell we defined a new coordinate system by empirically finding the unit vector that maximized the projection of the CS-induced trajectory change at peak velocity. For example, the right side of Fig. 3D shows a saccade’s trajectory in velocity space with and without a preceding CS. We can compare the two trajectories at peak speed and compute the difference in the two velocity vectors. Let us label this difference, as measured across all saccades, as the vector that maximizes the CS-induced disturbance.

For each P-cell, we compared the maximum vector with the CS-on vector that we had measured for that cell. The distribution of the angular difference is shown in Fig. 3D. The distribution is not uniformed as would be expected based on bootstrap analysis (Fig. 3D, shaded area shows 95% CI), but rather shows a peak for near CS-on angles and a trough for near CS+180 angles (50% of cells have maximum vector within 60 deg centered at CS-on whereas only 20% of cells have maximum vector within 60 deg centered at CS+180, p<0.01 as measure by a bootstrapping procedure). Thus, the CS-on vector was an unbiased estimator of the optimal vector.

However, for many P-cells, the maximum vector was 90° or more apart from CS-on. Indeed, when the difference in the with- and without-CS saccades were projected onto the maximum vector of each P-cell, the results produced a larger trajectory changed (Fig. 3E, maximal trace) than what we had measured when the data were projected onto CS-on (Fig. 2A). Is the maximum vector a better coordinate system with which to measure the CS-induced change in the saccade?

To answer this question, we performed the same bootstrap analysis as in Fig. 2A. That is, we separated the saccades randomly into two groups (i.e., not based on whether there was a CS or not). The bootstrap analysis produced a 95% confidence interval, effectively estimating the size of the noise associated with the maximum vector (Fig. 3E, 95% CI). For the maximum vector, the distance of the effect size to noise was much smaller (1.5 deg/s at peak velocity) than the distance as measured for the CS-on vector (4.5 deg/s at peak velocity). Thus, the CS-on vector was both an unbiased and a robust estimate of the direction in which the CS had perturbed the saccade.

### Is the CS causing a perturbation, or simply correlated with a change in the saccade’s trajectory?

Our interpretation has been that the arrival of a spike from the inferior olive produced a CS that briefly suppressed the SSs that the P-cell would normally produce during the saccade, and this culminated in commands that pulled the eyes toward direction CS-on of that P-cell. However, it is also possible that in those trials in which there was a CS before saccade onset, the subject happened to plan its movement to a position that was slightly biased toward direction CS-on of the recorded P-cell. Is there a way to distinguish between the ‘perturbation’ and the ‘planning’ viewpoints? To explore this question, we first asked whether the CS activity before saccade onset carried information about the direction and amplitude of the impending movement, and then used that information to make predictions that could establish whether CS was causally affecting the movement, or merely correlated with the change.

In principle, the CS activity should carry information about the ensuing saccade because the olivary input to the oculomotor vermis transmits in part the activity of neurons in the superior colliculus (16–18). Collicular neurons, particularly those in the intermediate layers, can have a burst of activity in response to a visual stimulus, and then a second burst to instruct a gaze change toward that stimulus (19–21). Indeed, following presentation of the target, the CS firing rates (in direction CS-on) exhibited two peaks (Fig 1D): the first peak appeared to follow the onset of the visual target, while the second peak appeared to occur right before saccade onset, both with a preference to the same region of the visual space (the CS-on direction). This raised the possibility that the first CS peak was in response to the location of the visual stimulus, reflecting collicular encoding of the target in visual space, while the second peak was an encoding of the motor related collicular activity that instructed the movement to that location. Of course, in most cases the externally instructed goal and the internally selected movement were the same. However, there were instances in which the animal was shown a visual target but chose to ignore it, instead making a saccade elsewhere. We termed these *incorrect saccades* (Fig. 4A).

**Figure 4.**
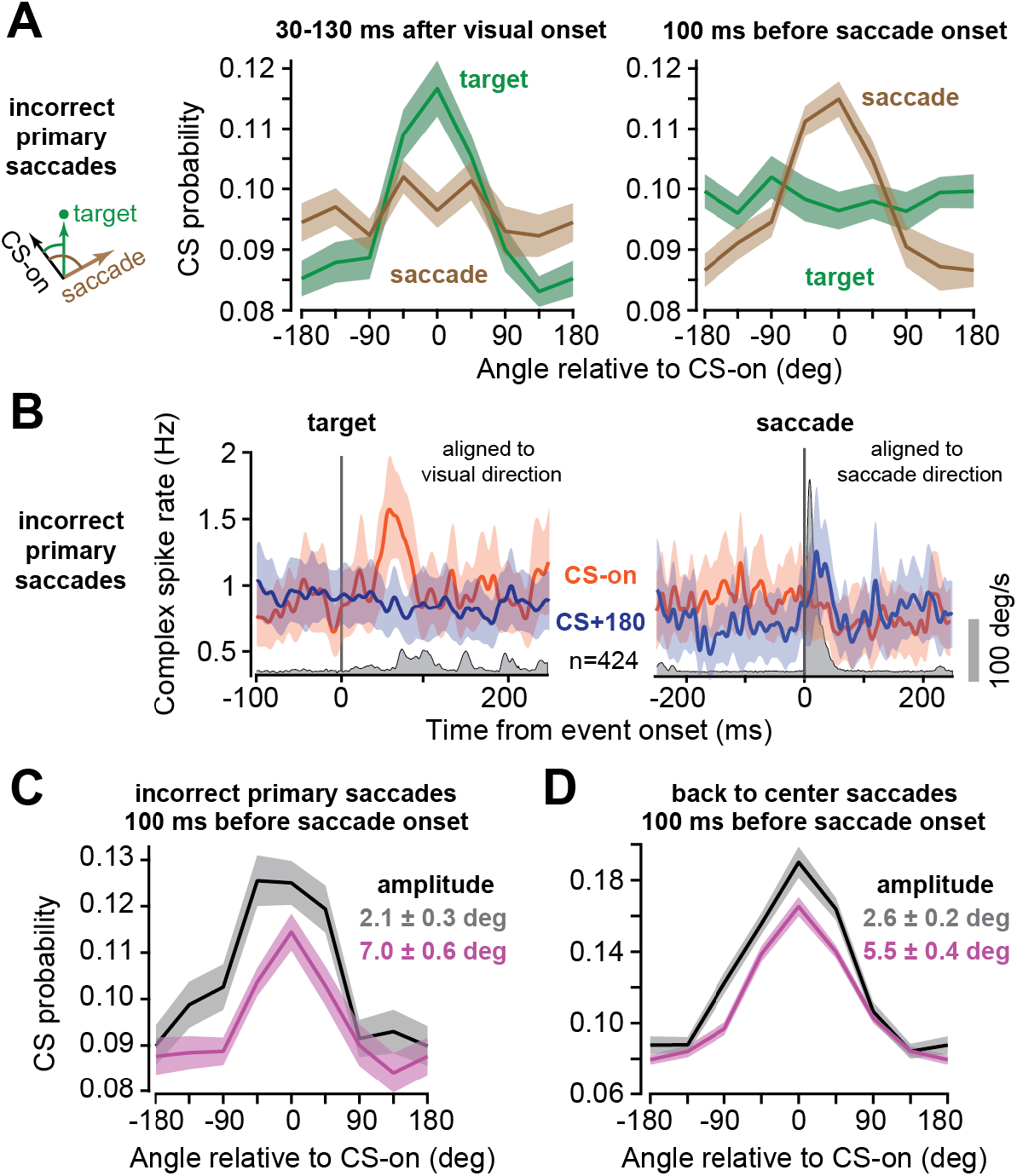
Complex spikes convey information about both the target location, and the intended movement. **A**. In the incorrect saccades, the subject was presented with a visual stimulus, but chose not to make a saccade toward it. At 30-130 ms following stimulus onset, the CS response was tuned to the direction of the target, not the direction of the eventual saccade. At 100 ms before saccade onset, the CS response was tuned to the direction of the upcoming saccade, not the direction of the target. **B**. Average firing rates for CS-on (which includes CS±45) versus CS+180 (which includes CS±135) showing higher response for CS-on visual target and higher response prior to movements in direction CS-on. **C-D.** For incorrect saccades, as well as back to center saccades, regardless of saccade direction the CS response at 100 ms before saccade onset was larger if the upcoming movement had a smaller amplitude. Error bars are SEM.

For the incorrect saccades, the CS response that followed the onset of the visual target (30-130 ms period) showed an early peak and a tuning that was aligned to the direction of the target with respect to CS-on (Fig. 4A-B, left subplots). This suggests that the early CS response encoded the stimulus location, despite the fact that the eventual movement was elsewhere. Indeed, there was no tuning of the CS response when aligned to the direction of the actual saccade (Fig. 4A, left subplot). That is, the initial CS response encoded the visual stimulus, but not the saccade that would eventually be made.

However, for the incorrect saccades, the CS response that preceded the movement (−100 to 0 ms) was no longer tuned to the direction of the instructed target (Fig. 4A, right subplot). Rather, it showed an elevated response (though not a distinct peak response) and tuning to the actual saccade direction with respect to CS-on (Fig. 4A-B, right subplots). This tuning is unlikely to result from a perturbation of the saccades to move in direction CS-on which shifts the CS tuning, as simulations we performed showed that the magnitude of the observed tuning (which shows >10% deviation from baseline) is too large to result from perturbations on the order of 1% (Fig. S2A-B). Thus, these results suggested that the CS activity following target onset reflected the location of the instructed stimulus and not the eventual movement. In contrast, the CS activity before saccade onset encoded the direction of the movement and not the visual stimulus that instructed it. (Note that for incorrect saccades the time between visual target onset and saccade onset had a mean of 350 ms and standard deviation of 360 ms, whereas for primary saccades it was 195±93ms).

Critically, we found that the CS activity before saccade onset reflected not just the direction of the ensuing saccade, but also grossly the amplitude of that movement. For all directions of movement, the CS probability preceding small amplitude saccades (defined as saccades below 3.5 deg, see Methods) increased by about 10% relative to larger amplitude saccades (Fig. 4C). We found similar results for back-to-center saccades (Fig. 4D). These results are consistent with prior findings that low amplitude visual targets have a higher probability of eliciting a complex spike than high amplitude visual targets (22, 23).

Together, the results suggested that there were two different kinds of information encoded by the CS firing rates: an early signal that encoded the visual event, and a later signal that encoded the internally chosen movement direction. The early and the late CS rates showed a preference to the same region of the visual space (the CS-on direction) but differed if the subject chose to move to a location that was not instructed. Thus, the CS rates before saccade onset conveyed information about the direction and amplitude of the movement that the subject planned to make.

We are now in a position to formulate the “planning” hypothesis. The CS rate in each P-cell will be greater when the ensuing movement direction is closer to the CS-on direction of that P-cell. Moreover, the CS rates will be greater when the movement has a smaller amplitude (Fig. 4A). In this framework, the CS does not perturb the saccade. Rather, it merely conveys information about the upcoming movement.

The planning hypothesis makes a critical prediction: regardless of movement direction, presence of a CS should signal a smaller amplitude movement (Fig. 5A, left panel, red marks). In contrast, the perturbation hypothesis makes a different prediction: because the presence of a CS will always pull the eyes in direction CS-on, the effect of the CS will be direction dependent (Fig. 5A, left panel, black dots). Notably, if the target is in direction CS-on and there is a CS, then the saccade will have a slightly larger amplitude (with respect to no CS). However, if the target is in direction CS+180 and there is a CS, then the effect of the CS will be to slow the saccade, thus producing a slightly smaller amplitude.

**Figure 5.**
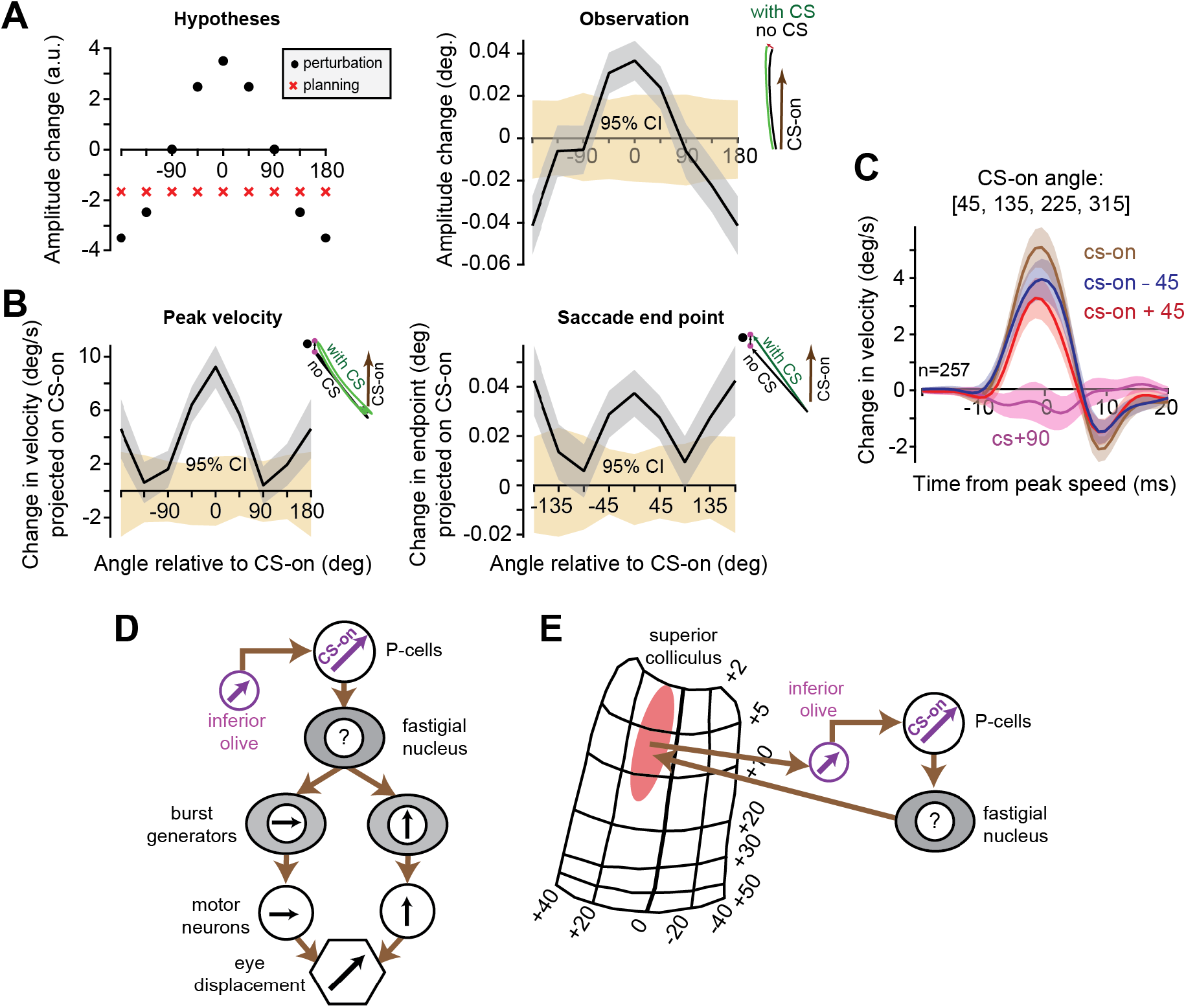
Comparing the perturbation hypothesis with the planning hypothesis. **A**. Left: predictions of the two hypotheses. Right: the measured data. Presence of a CS had a direction dependent effect on saccade amplitude, consistent with the predictions of the perturbation hypothesis. **B.** Projection of the CS induced changes in peak velocity (left subplot) and in final saccade position (right subplot) on the CS-on vector. Perturbation of movements in directions CS-on and CS+180 are larger than perturbation of movements in directions CS±90. **C.** For P-cells whose CS-on angle is along the diagonal axes the perturbation is strongest when it is projected on the CS-on vector, weaker when projected on the CS-on±45 vectors. **D-E.** Hypothetical diagrams showing the sensorimotor map of a P-cell whose CS-on is at 45 deg. If the downstream nucleus neuron projects onto burst generators, then that projection must be to both vertical and horizontal burst generators. If the nucleus neuron projects to the superior colliculus, then that projection should be to the same region that provides CS information to the P-cell. The tuning of fastigial nucleus is yet to be explored. Error bars are SEM or 95% CI.

We measured saccade amplitudes for each target in the condition when there was a preceding CS vs. when there was no CS. Indeed, when the target was in direction CS-on, the presence of a CS was followed by a slightly longer saccade. In contrast, if the target was in direction CS+180, then the presence of a CS was followed by a slightly shorter saccade (Fig. 5A, right panel; amplitude difference between CS-on and CS+180 is ∼0.08 deg, both values are with p<0.01 as measure by a bootstrapping procedure). These results are consistent with the differences in peak speed between CS-on and CS+180 movements (Fig. 2D) and agree with the predictions of the perturbation hypothesis but not the planning hypothesis.

A further hint is that for movements in directions CS-on and CS-180, the CS induced changes are larger than for movements that are orthogonal to the CS-on angle (CS±90). To show this, we computed for all 8 movement directions the CS induced changes in the maximum velocity vector (Fig. 5B left subplot) and saccade endpoint vector (Fig. 5B right subplot) by subtracting for each cell the average without-CS-saccades vector from the average with-CS-saccades vector. We then projected the resulting vectors on the CS-on vector and averaged across all cells. While the resulting scalar value is positive across all movement directions (suggesting that in all movement directions there is a perturbation in direction CS-on), the magnitude of the perturbation is not equal across movement directions (movements in directions CS±90 are within 95% CI of random splitting of the data, whereas movements in direction CS-on and CS+180 have p<0.01; movements in directions CS±45 also have, on average, larger perturbation than movements in directions CS±135). As P-cells are unlikely to be involved in movements that are orthogonal to their CS-on angle, a pause in their SS had a smaller perturbation effect on those movements. On the other hand, if the change in saccade trajectory following a CS reflected the higher probability of movements that are closer to the CS-on direction, then it is hard to understand why the change is larger for CS-off movements, which have the lowest CS probability, than for orthogonal movements.

In summary, the CS rates before saccade onset carried information about the direction and amplitude of the upcoming movement. This raised the possibility that the presence of a CS did not disrupt the movement, but only signaled that the movement would be closer to the preferred direction of the P-cell. However, the planning hypothesis could be dissociated from the perturbation hypothesis by measuring the direction-dependent amplitudes of the saccades. We found that the presence of a CS was followed by a saccade that was longer when directed to a target at direction CS-on, but shorter when directed to a target at direction CS+180. This result was consistent with the perturbation hypothesis, suggesting that the CS disrupted the movement, pulling the eyes toward direction CS-on of the P-cell that had produced the complex spike.

### Is the output of the P-cell aligned with muscle coordinates or sensory coordinates?

Downstream of the cerebellum and the superior colliculus are the burst generator neurons that activate the motor neurons and drive the eye muscles. The tuning functions of both the burst generator neurons and the motor neurons are along the vertical and the horizontal axes (Fig. 5D) (24–28), reflecting the axes of eye muscles. CS tuning of P-cells, on the other hand, is aligned to a polar coordinate system, similar to neurons in the superior colliculus, reflecting the visual space about the fovea. This difference between the sensory and motor spaces allowed us to ask a new question: do P-cells with CS-on along the diagonal axes, which are not aligned with burst generator neurons, drive movement in the direction of their CS-on, or along the tuning of the downstream burst generator neurons (i.e., in the CS-on±45 direction)? To answer this, we projected the velocity difference between with- and without-CS saccades on the CS-on, CS-on+90 and CS-on±45 vectors but just for cells whose CS-on was along the diagonal axes. We found that, on average, the strongest perturbation was in direction CS-on (Fig. 5C). Furthermore, we repeated the analysis of Fig. 2D and Fig. 5A analyzing differences in peek speed and saccade amplitude between with- and without-CS saccades for the different direction movements, but now just for cells whose CS-on is along the diagonal axes. We again found that the largest difference in speed and saccade amplitude was in direction CS-on while the smallest difference in speed and saccade amplitude was in direction CS-180 (Fig. S2C).

In summary, when we focused exclusively on the P-cells that had a CS-on along the diagonal axis, the effect of the CS was to pull the eyes along the same diagonal axis, not along the horizontal or vertical axes. This implies that on average, the downstream effect of a P-cell is to drive a movement in the direction specified by the inferior olive, even when that direction is not aligned with the coordinates of any single muscle.

## Discussion

It is difficult to infer the relationship between the simple spikes that a P-cell is generating during a movement, and its downstream effects on behavior. Here, we found that the encoding of information in the climbing fiber input that a P-cell receives provides a critical clue. On the one hand, a climbing fiber input to a P-cell in the oculomotor vermis encodes a specific region of the visual space, producing a CS that signals that a visual event has taken place around that location. On the other hand, when the CS occurs near saccade onset, it suppresses the SSs and alters the saccade’s trajectory, pulling the eyes toward the same region of the visual space that was preferentially encoded by the climbing fiber input. This implies that the sensorimotor map of a P-cell is defined by its climbing fiber input. When the simple spikes are suppressed, this signals the downstream motor structures to produce commands that will pull the eyes in direction CS-on.

In both macaques and marmosets, when a saccade is made in direction CS+180, the population SS rate initially exhibits a burst but then falls below baseline just before the movement begins to decelerate (12, 14). From the current results we infer that the SS suppression is signaling the downstream motor structures to produce forces in direction CS-on, which would oppose the ongoing movement, thus bringing it to a stop. Moreover, because P-cells that are located on one side of the vermis tend to have a CS-on direction that is biased toward the contralateral side (14), the results suggest an explanation as to why a unilateral lesion of the cerebellum produces hypermetric saccades in the contralateral direction (29–32).

We interpreted the effects that the CS had on behavior to be due to SS suppression, which results in the disinhibition of the nucleus neuron, making it burst (33). However, there are other possibilities. For example, the CS itself is a spike that is transmitted down the P-cell’s axon (34, 35). Notably, the downstream effect on the nucleus neuron is inhibition, particularly when the CSs are synchronized among the P-cells (36–39). The increased inhibition of the nucleus, for example via stimulation of P-cells, tends to drive the eyes opposite to the direction of CS-on (40). Another possibility is that an inferior-olive neuron that sends a climbing fiber to a P-cell also sends collaterals to the cerebellar nucleus neuron that receives input from the same P-cell (33, 41, 42). If this is indeed the case (38), then the olive can directly activate the nucleus, potentially bypassing the need to suppress the P-cell’s simple spikes. However, this alternate scenario may be less effective than SS suppression of the parent P-cell because olivary inputs to nucleus neurons tend to be weak, at least in the adult mouse (43, 44).

A central idea in cerebellar research is that the CS is a teaching signal for the P-cell (45–48). Indeed, when a saccade ends and the target is not on the fovea, the resulting visual error modulates CS probability (12, 22), inducing learning which is then expressed through modification of the SSs during the subsequent movement (13). Our results here imply that if the visual error produced an increase in the CS probability of a P-cell, then the subsequent reduction in that P-cell’s SSs has a specific meaning: the motor system is instructed to pull the eyes in the direction of the error vector, thus reducing that error.

However, as noted in many experiments, CSs occur not just in response to error, but also before and during a movement (15, 49–53). Critically, CSs occur before movement onset even when there are no sensory events that triggered that movement (54). Here we found that CSs that occurred before a saccade carried information about the internally selected direction of the movement, not the visually instructed direction. What might be the purpose of CSs that do not signal an external event or an error?

As pointed out by Bouvier et al. (55), a general learning system cannot rely solely on a teacher to provide it with information regarding how to correct an error. Indeed, for many cerebellar-dependent behaviors, the cerebellum is not afforded the luxury of a teacher (56). In theory, an effective solution for the cerebellum would be to have a way to stochastically perturb the movement, produce a change in behavior, then evaluate that change to determine its utility (55). Our results provide evidence that at least part of this mechanism may be employed by the cerebellum: CSs that occur near the onset of a saccade perturb that movement. However, the effect size that we observed was small, altering saccade trajectory by adding a vector that was 1% of the peak velocity of the movement. This effect size was tripled when the CSs were synchronized. Thus, the olivary input to the cerebellum can stochastically alter movements, particularly by synchronizing this input among the P-cells. In theory, the purpose of this input may be to induce stochastic learning without a teacher.

While the CS-induced perturbation pulled the eyes in direction CS-on, the resulting displacement appeared to be partially corrected by subsequent commands near the end of the saccade (Fig. 2B). Thus, there appears to be a mechanism in place to monitor the ongoing movement and at least partially correct for the internally generated perturbation (57, 58). Indeed, earlier we found that when saccades were disrupted via transcranial magnetic stimulation, the brain maintained the ability to correct the movement and steer the eyes to the target (59). It will be intriguing to measure mossy fiber input and the activities of interneurons in the cerebellar cortex following a CS-induced perturbation and ask whether this circuitry monitors and then partially compensates for the perturbed movement.

Our results were obtained from the oculomotor vermis, a region that is principally concerned with the motion of the eyes. The eyes have a small mass that is more easily perturbed than other parts of the motor system, raising the question of whether our results can generalize to other parts of the cerebellum. An important clue is the work of Streng et al. (60) who trained macaques in a reaching task and recorded from the lateral regions of lobule IV-VI. They found that the occurrence of a CS during a reach was correlated with a subsequent motion of the hand, thus inferring that the CS was not reporting an error, but rather causing a change in the reaching movement. Their inference is consistent with our observations in the vermis, suggesting that across various regions, the olive has the means to perturb movements. Another interpretation, however, is that a CS reports the plan to make a movement, as we have shown here for saccade (the ‘planning’ signal). A critical question for future research is whether these perturbations are part of the stochastic gradient descent toolbox that may be employed by the olive to teach the cerebellum (55).

Our results suggest that the sensory representation of space in the climbing fiber can be transformed by a P-cell into a movement in the same direction. How might this transformation take place? One possibility is in the oculomotor vermis, a P-cell connects to a fastigial neuron that projects to the superior colliculus region (61) that forms the input to the olivary cell that innervates the same P-cell (Fig. 5E). In this way, all P-cells, including those that have a diagonal CS-on, can influence actions that match their olivary input. Another possibility is that the nucleus neuron projects to both horizontal and vertical burst generators in such a way that it can produce movements along a direction that matches the CS-on vector of the parent P-cell (Fig. 5D).

While P-cells in the oculomotor vermis have a CS tuning that reflects the visual map of the colliculus, in the flocculus the CS tuning is thought to reflect the motor map of the muscles. In the flocculus, the SS response is aligned to a Cartesian coordinate system with cells tuned mostly to right/left and up/down smooth-pursuit movements (62). CS tuning for these cells is inversely aligned with the SS tuning (63). What may underlie the difference in the CS tuning of these two cerebellar structures? The flocculus P-cells are one synapse away from eye motor neurons (with the vestibular nuclei separating them), whereas vermis P-cells are 2 or 3 synapses away (with the fastigial nucleus, superior colliculus, and burst generator neurons separating them). Thus, perhaps the added neuronal layers from the vermis to motoneurons enable the transformation from the vermis polar coordinate system to the motor neuron Cartesian coordinate system.

Finally, our results that CSs can produce movements sheds new light for certain diseases of the inferior olive. In oculopalatal tremor (OPT), during fixation there is smooth, low frequency oscillation of the eyes and the palate (64). OPT is believed to occur when a lesion or disease disrupts the inhibitory inputs from the cerebellar nuclei to the inferior olive. This loss of inhibition leads to hypertrophy of the inferior olive, and the cells develop abnormally strong soma-somatic gap junctions. Our results suggest that the motor symptoms of OPT may be a result of excessive climbing fiber input to the cerebellum, causing unwanted movements.

## Methods

Data were collected from two marmosets over the course of 2.5 years (*Callithrix Jacchus*, male and female, 350-370 g, subjects M and R, 6 years old at start of recordings). The marmosets were born and raised in a colony that Prof. Xiaoqin Wang has maintained at the Johns Hopkins School of Medicine since 1996. The procedures on the marmosets were approved by the Johns Hopkins University Animal Care and Use Committee in compliance with the guidelines of the United States National Institutes of Health.

### Data acquisition

Following recovery from head-post implantation surgery, the animals were trained to make saccades to visual targets and rewarded with a mixture of applesauce and lab diet (65). Visual targets were presented on an LCD screen (Curved MSI 32” 144 Hz - model AG32CQ) while binocular eye movements were tracked using an EyeLink-1000 eye tracking system (SR Research, USA). Timing of target presentations on the video screen was measured using a photo diode. We performed MRI and CT imaging on each animal and used the imaging data to design an alignment system that defined trajectories from the burr hole to various locations in the cerebellar vermis (65), including points in lobule VI and VII. We used a piezoelectric, high precision micro drive (0.5 micron resolution) with an integrated absolute encoder (M3-LA-3.4-15 New Scale Technologies) to advance the electrode. We recorded from the cerebellum using quartz insulated 4 fiber (tetrode) or 7 fiber (heptode) metal core (platinum/tungsten 95/05) electrodes (Thomas Recording), and 64 channel checkerboard or linear high density silicon probes (M1 and M2 probes, Cambridge Neurotech). We connected each electrode to a 32 or 64 channel head stage amplifier and digitizer (RHD2132 and RHD2164, Intan Technologies, USA), and then connected the head stage to a communication system (RHD2000 Evaluation Board, Intan Technologies, USA). Data were sampled at 30 kHz and band-pass filtered (2.5 - 7.6 kHz). We used OpenEphys (66) for electrophysiology data acquisition, and then used P-sort (67) to identify the simple and complex spikes in the heptodes and tetrodes recordings, and Kilosort and Phi (68) to identify the spikes for the silicon probes. The simple spikes in a part of this data set were analyzed and reported previously (14).

### Behavioral protocol

Each trial began with fixation of a center target for 200 ms, after which a primary target (0.5×0.5 deg square) appeared at one of 8 randomly selected directions at 5-6.5 deg (50 cells were recorded during sessions with 4 random directions). The onset of the primary target coincided with presentation of a distinct tone. As the subject made a saccade to this primary target, that target was erased, and a secondary target was presented at a displacement of 2-2.5 deg, also at one of 8 randomly selected directions (4 directions for 50 cells). Following 200 ms fixation of the final target, reward was presented with a distinct tone, and the center target was displayed.

### Data analysis

All saccades, regardless of whether they were instructed by presentation of a visual target or not, were identified in the behavioral data using a velocity threshold. CS and SS rates were smoothed with a 20 ms Gaussian kernel.

#### Computing the CS-on vector

CS directional tuning was computed by measuring the CS firing rates following target onset as a function of target angle with respect to the actual position of the eyes. We counted the number of CS after target onset up to saccade onset or a fixed 200 ms window, whichever happened first. Dividing the spike count by the duration of time resulted in the CS firing rate. We then weighed the vectors pointing to the target by their firing rates and summed them. We defined the resulting angle as the CS-on angel.

#### Rounding CS-on angle

We rounded the CS-on to the nearest 45 deg angle and thus classified cells as having the same CS-on if their rounded CS-on angle was the same. Thus, for the population analysis of Fig. 2B this criterion produced 35 sessions that we had two P-cells, 6 sessions with three P-cells, 8 sessions with four P-cells, 5 sessions with five P-cells, 2 sessions with six P-cells, 2 sessions with seven P-cells, and 1 session with eight P-cells that had the same CS-on (total of 179 P-cells)

#### Rounding movement directions

Similarly, for defining movement directions as belonging to one of 8 directions we rounded the movements to the nearest 45 deg angle. We defined a movement direction based on the starting point and target end point. However, when analyzing distributions based on actual saccade directions (Fig. 4) we defined movement direction based on the starting point and actual end point.

#### Using weighted sum

To account for the fact that some cells were recorded for much longer period than other cells we used weighted sum across cells (weighted by the number of CSs in the relevant time window) for mean and SEM analyses. Similarly, to account for the fact that within cells some movement directions had many more CSs than other directions we used weighted sum when averaging across directions. Thus, the bootstrap analysis randomly separated the movements, but maintained the ratio of with-CS and without-CS saccades for each direction and each cell.

#### Vector projection

Projection of a vector implies taking the dot product of the analyzed vector with a unit vector defining the CS-on, CS+90, CS±45 or the maximal vector.

#### Effect of saccade amplitude

For separating saccades based on amplitude we defined 3.5 deg amplitude to be the threshold as it was close to the median. But we tested with varying thresholds and found that our results were not sensitive to the chosen threshold.

#### Comparing planning vs. perturbation hypotheses

To construct the perturbation hypothesis of Fig. 5A, we added to each direction a vector in a defined CS-on direction. We then calculated the change in amplitude of each vector caused by the addition of the CS-on vector. For the ‘planning hypothesis’ we just plotted all to be below zero as probability for a CS was larger for low amplitude saccades.

#### Fig. S1B

We analyzed changes in saccade amplitude and projection of changes in saccade endpoint while normalizing for the differences in the saccade start point. We used the following to compute the CS induced change in saccade amplitude: 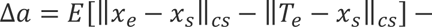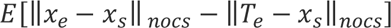, where *x*_s_ and *x_e_* are the start and endpoint of the saccade, and *T_e_* is the end target location. The symbol 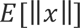 indicates the expected value of the L2 norm of the vector *x*. Effectively, we first subtracted the instructed amplitude from the actual amplitude for each saccade, and then averaged this quantity for with- and for without CS-saccades and then subtracted the value for without-CS-saccades from the with-CS saccades. We then averaged this quantity across cells to compute the effect of the CS on the deviation of the saccade amplitude from the instructed saccade amplitude. Similarly, to compute the projection of changes in final saccade point we measured the following:

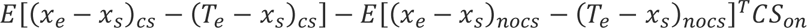, where *x^T^* indicates the transpose of the vector and *CS_on_* is the CS-on unit vector.

### Statistics

To estimate the statistical significance of the data we used a bootstrapping method in which we randomly split the data into two groups, a fake CS group and a fake no-CS group. Importantly, we maintained the numbers in the groups as in the real splitting of the data. We then ran the analysis exactly as in for the real data 1000 times (with different random seeds) and extracted a 95% and a 99% confidence interval (CI). The statistic p<0.01 and p<0.05 mean that the real data were outside the 99% and 95% CI, respectively.

## Acknowledgements

The work was supported by grants from the NIH (R01-EB028156, R01-NS078311), and the Office of Naval Research (N00014-15-1-2312). S.Z.M. was further supported by NSF NeuroNex award (1707398) and by the Gatsby and the Swartz Foundations.

## Author contributions

J.S.P., P.H., M.A.F., E.S., and R.S. conceived and performed experiments and pre-processed the data. S.Z.M. and R.S conceived the project, analyzed the data, made the figures and wrote the manuscript.

---

**Figure S1.**
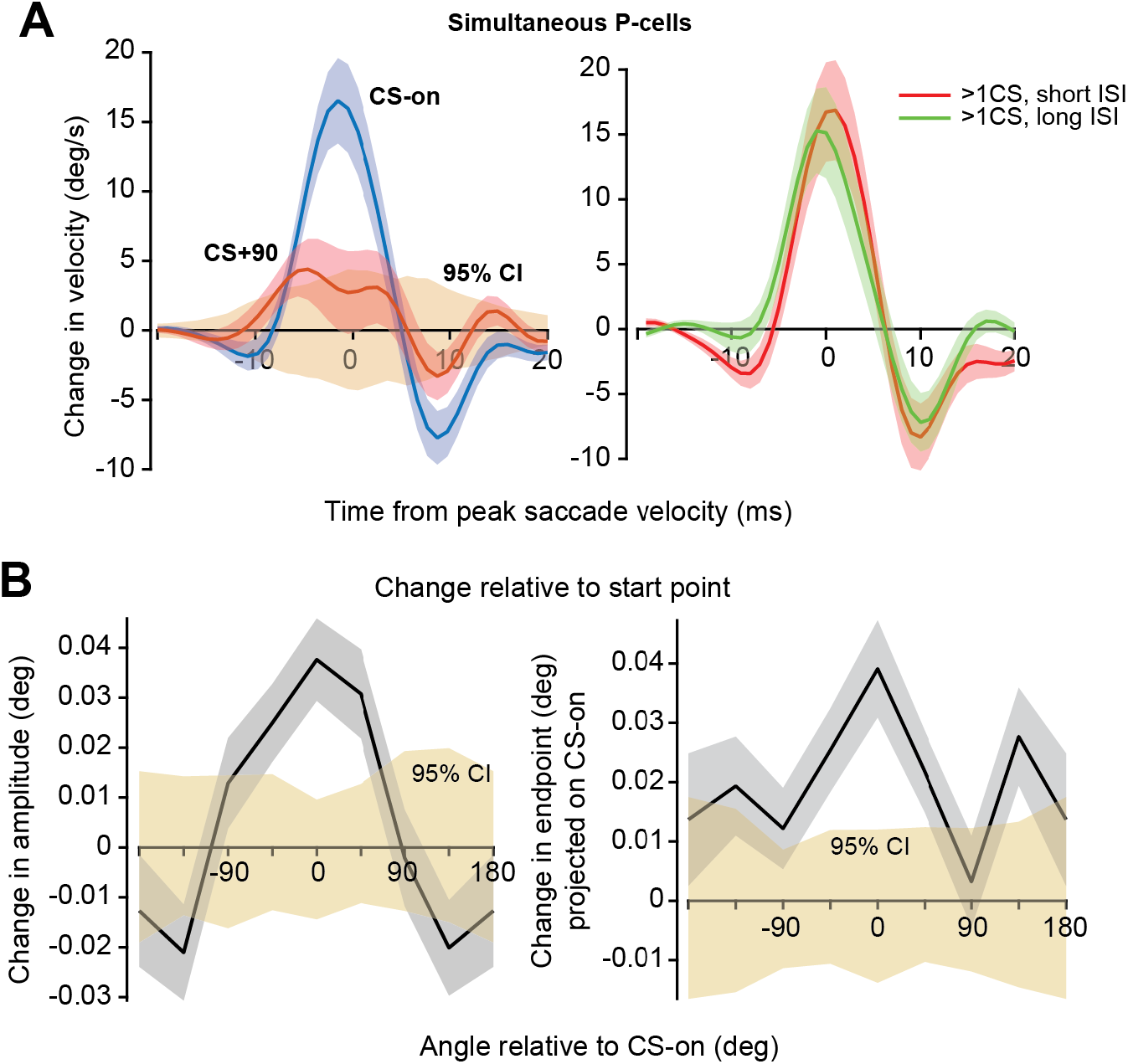
Simultaneous CSs produced larger perturbations, and the perturbations were robust after normalizing for the small variations in saccade start point. **A.** Left: similar to Fig. 2B, but here we added the 95% CI based on bootstrapping the random splitting of the data. Right: we divided the data based on the temporal distance between complex spikes, based on median of 15 ms. **B.** Similar to the analyses of Fig. 5A and Fig. 5B, but here we accounted for the difference in saccade start points by first subtracting the instructed saccade from the actual saccade (see Methods for details). Thus, here we analyzed the effect of a CS on the deviation of the actual saccade from the instructed saccade. Error bars are SEM and 95% CI.

**Figure S2.**
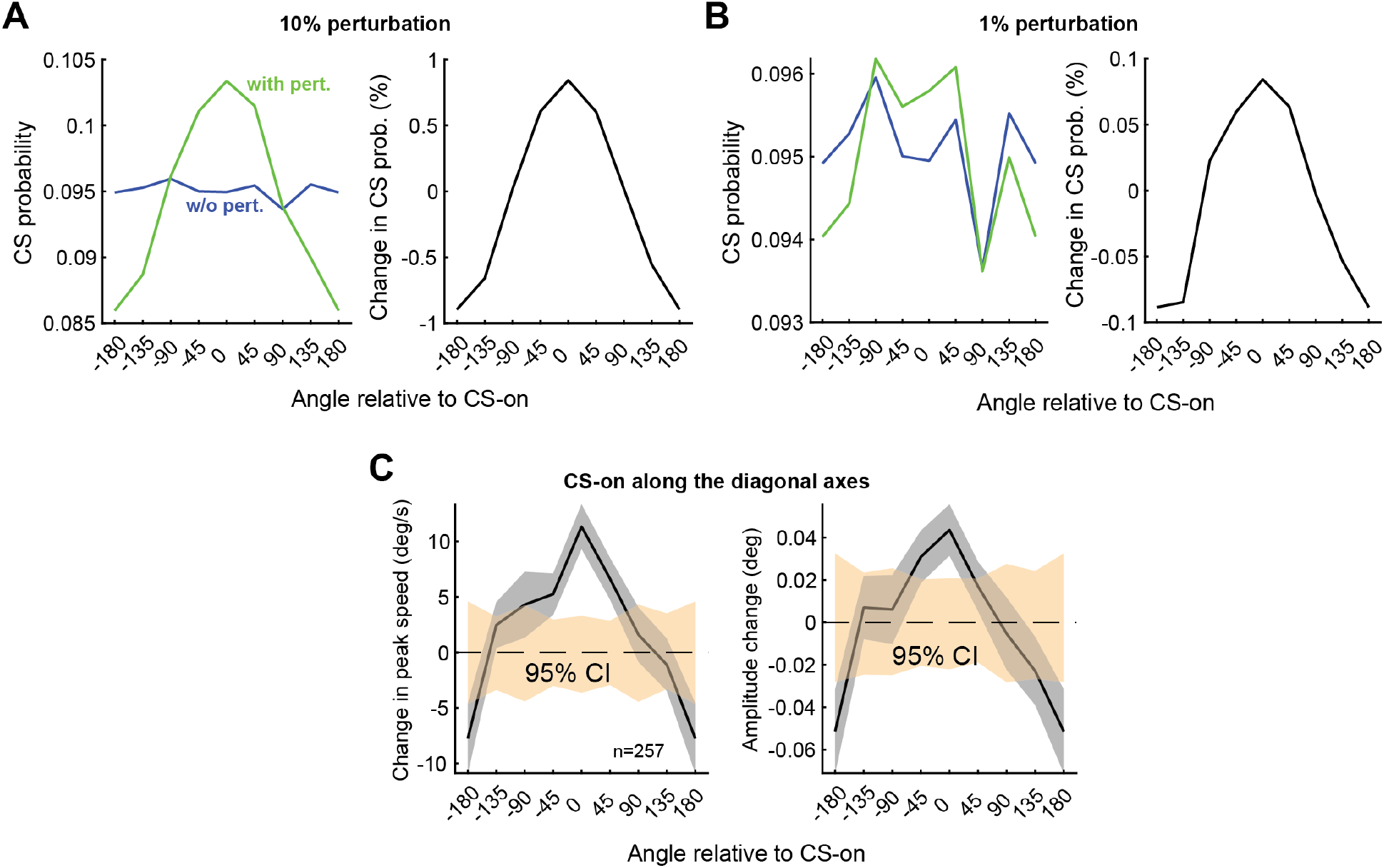
Simulation results suggest that the CS tuning of P-cells is not influenced significantly by the fact that the CS perturbs the saccade. Moreover, perturbation in direction CS-on best explains the effect of a CS even for cells in which the CS-on angle is along the diagonal axes. **A-B.** We simulated saccade angles along a uniform distribution, with the amplitude drawn from a Gaussian distribution with a mean of 5 deg and std of 0.7 deg. 10% of the saccades were randomly assigned to be with-CS-saccades. We then analyzed the CS tuning of the saccade to confirm that it was uniform across movement directions (left subplots ‘without perturbation’). We then added a perturbation condition (‘with perturbation’) in which we added a vector in direction CS-on to the end of all with-CS saccades. Next, we varied the magnitude of the perturbation vector to be large (10%), or small (1%, as in our measured data). Finally, we analyzed the CS probability as a function of saccade angles of the with- and without-CS saccades. To account for the noise in the simulation, the right subplot shows the change in the CS tuning caused by the perturbation (i.e., we computed CS probability of with-CS minus without-CS saccades). The magnitude of the change in the CS tuning was proportional to the magnitude of the perturbation: 10% perturbation caused a shift of ∼10% in the CS probability (for CS-on and CS+180) while 1% perturbation caused a shift of ∼1% in the CS probability. **C.** Same as Fig. 2D and Fig. 5A but here just for cells whose CS-on angle is along the diagonal axes. Error bars are SEM or 95% CI.

